# Gene expression in *Anolis sagrei* suggests conservation of patterning mechanisms drives formation of two high acuity areas

**DOI:** 10.64898/2026.11.03.764084

**Authors:** Heer N.V. Joisher, Isabella van der Weide, Bonnie K. Kircher, Richard R. Behringer, Constance Cepko

## Abstract

As humans we enjoy high acuity color vision. A specialized area in our retinas, the fovea, endows us with this ability. Some vertebrates, such as the *Anolis sagrei (brown* anole), have two such areas. Whether different species use conserved developmental mechanism(s), and whether the same mechanism is used in one organism to pattern both foveae, are herein addressed in the brown anole. We observed that the expression of canonical axis patterning genes is conserved in the anole retina. Retinoic acid and Fgf signaling pathways were found to define two discrete domains corresponding to emerging foveae. Comparative spatial analyses of early gene expression patterns among anole, chick, and human revealed conserved patterns in the anlage of the high acuity areas of each species. Variation in foveal number could arise through spatial redeployment of conserved patterning programs.

## Main Text

Spatial organization is a central feature of tissue development, ensuring that distinct functional regions emerge at defined positions within a developmental field (Dutting and Thanos, 1995; Echelard et al., 1995; Crossley et al., 1996). The vertebrate retina provides a clear example of this principle, organized not only in laminae comprising cellular layers, but also in specialized topographic domains that directly shape visual performance (Matsuno et al., 1992; McLaughlin and O’Leary, 2005). Prominent among these specializations are high-acuity areas (HAAs), which are characterized by differences in photoreceptor composition, including increased density of cone photoreceptors and absence of rod photoreceptors, as well as elevated ganglion cell density. In addition, in some species, displacement of retinal cell layers creates a pit (fovea) (Hendrickson, 1992; Hendrickson and Yuodelis, 1984). HAAs exhibit variation in number, position, and structural organization, reflecting selection for visual system performance across species. How these specialized regions are established is an active area of investigation. In primates, high-acuity vision is enabled by a single fovea, positioned slightly temporal to the centrally localized optic nerve head, and is characterized by cone subtype enrichment, absence of rods, specialized circuitry, and a retinal pit (Hendrickson, 1992). In other vertebrates, the localization is slightly different. Birds typically possess a high-acuity area positioned slightly nasal to the optic nerve head. It has a specialized cellular composition, similar to that of humans. The central HAA of birds can exist with (e.g. Zebrafinch) or without (e.g. Chicken) a foveal pit, and some foveated species (e.g. eagle) feature a second, temporal fovea (da Silva and Cepko, 2017; Portier et al.,2017; Sugiyama et al., 2020). Comparable multi-foveate configurations are also observed in reptiles; among these, the brown anole (*A. sagrei)* has two spatially distinct foveae, a deep central fovea and a shallower temporal fovea (Rasys et al.,2021; Sannan et al., 2018; Rasys et al., 2024; Rasys et al., 2025; Wahle et al., 2023) (Figure 1A). Despite the importance of these areas to vision, the developmental mechanisms that determine the number and spatial positioning of high-acuity regions remain poorly understood.

**Figure 1.**
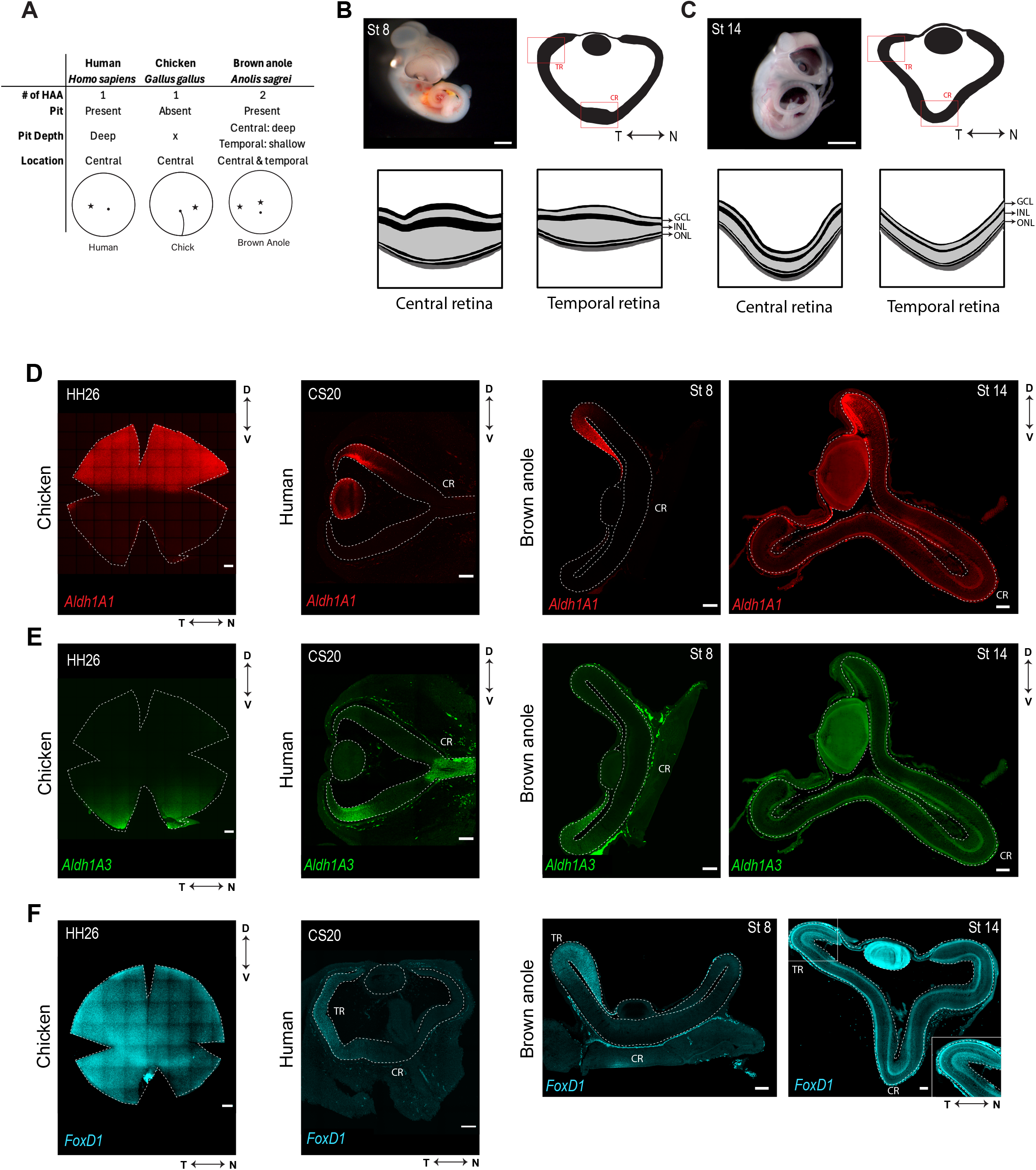
Comparative organization of retinal high acuity areas across human, chick, and brown anole retina. **(A)** Summary of high acuity area (HAA) features in human (*Homo sapiens*), chicken (*Gallus gallus*), and brown anole (*Anolis sagrei*). Schematic retinal diagrams below each column indicate the approximate position of the HAA/foveal specializations as seen from the back of the eye ball, with stars marking HAA locations and dots/lines marking the optic nerve head for anole/human and optic fissure for chicken. **(B-C)** Whole embryos and schematic retinal cross-sections during early stages of retinal development in *Anolis sagrei*. (B) Stage 8 (St8) embryo and schematic NT cross-section illustrating early retinal organization. (C) Stage 14 (St14) embryo and schematic NT cross-sections showing developing central and temporal retinal regions. Insets show higher-magnification views of central and temporal retinal domains corresponding to future high-acuity regions. Cross section schematics adapted from Rasys at al., 2025. **(D-F)** Comparative gene expression patterns across developing chick, human and brown anole retinas. For each panel, images are arranged from left to right as follows: chick (HH26 or HH28 as indicated); human (CS20); brown anole St8; brown anole St14. (D) *Aldh1a1*; (E) *Aldh1a3*; (F) *FoxD1*. D, dorsal; V, ventral; N, nasal; T, temporal; CR, Central Retina; TR, Temporal Retina. Scale bar – (A) 5mm; (B-C) 2mm; (D-F): Chick (200 μm), Human (100 μm), Brown Anole (100 μm). All patterns were observed across at least 2 biological replicates (N > or = 2).

Patterned gene expression in the developing retina establishes positional information along the dorsoventral (DV) and nasotemporal (NT) axes, providing a coordinate system for regional specialization (Sakuta et al., 2006, Takahashi et al., 2009). These spatial patterns arise early in eye development and are known to define domains of retinal identity that guide later structural and functional organization. Components of the retinoic acid (RA), fibroblast growth factor (Fgf), and bone morphogenetic protein (Bmp) pathways were first discovered to be associated with the developing HAA in chicks, and later also found in human embryos (da Silva and Cepko, 2017; Joisher et al., 2025). The circumscribed patterns suggest that these localized domains contribute to the development of the HAA and it’s positioning, and indeed functional perturbations of RA and Fgf showed that they do (da Silva and Cepko, 2017). However, it is unknown whether these same patterning systems have a conserved expression pattern in other vertebrate lineages, outside of mammals and birds. In addition, it was unknown whether a species with more than one HAA uses the same or different mechanisms to localize the different HAAs. The presence of two spatially distinct foveae in brown anoles provided an opportunity to address both of these questions.

We examined two developmental stages of the brown anole retina spanning early retinal patterning and foveal morphogenesis (Figure 1B and 1C). At an early stage, stage 8 (St8), localized retinal thickening marks the developing central and temporal foveal regions, where early lamination and sparse postmitotic cells are already evident, while much of the remaining retina appears to be comprised of mitotic retinal cells only. By stage 14 (St14), the developing foveal regions protrude outward and retinal lamination is more uniformly established throughout the tissue (Rasys et al., 2025). To determine whether early retinal axes are conserved in the brown anole, we examined the expression domains of canonical DV and NT patterning genes. In all species examined so far, including chicks, humans, mice, and zebrafish, retinoic acid synthesizing enzymes define complementary DV domains. *Aldh1a1 is* expressed dorsally in chick, human, and mouse retinas and *Aldh1a3* ventrally in chick, human, mouse and zebrafish retinas, typically with relatively broad and diffuse boundaries (Figure 1D and 1E) (Fan et al., 2003; Pittlik et al., 2008; da Silva and Cepko, 2017). In the brown anole, *Aldh1a1* was expressed in the dorsal retina at both St8 and St14. At St8, expression was observed broadly throughout the thickness of the retina, whereas by St14 it became more restricted to differentiated layers, specifically the ganglion cell layer and inner nuclear layer (Figure 1D). In contrast, *Aldh1a3* was detected ventrally at St8 at lower levels than *Aldh1a1* and was not detectable at St14 (Figure 1E). Overall, these DV expression domains appear restricted to the more dorsal and more ventral domains in the brown anole retina relative to those patterns of the human and chicken retinas.

Along the NT axis, *FoxD1* and *FoxG1* mark temporal and nasal identity, respectively, and exhibit complementary expression with relatively diffuse boundaries in the chick retina (Figure 1F, Supplementary Figure 1A) (Sakuta et al., 2006, Takahashi et al., 2009, Joisher et al., 2025). In the brown anole, *FoxD1* was enriched in the temporal retina at St8 and persisted at low levels at St14, in a more restricted domain (Figure 1F). *FoxG1* showed weak nasal expression at St8 and was not detected at the later stage (Supplementary Figure 1B). In both cases, expression domains were tightly restricted to temporal-most and nasal-most regions, respectively, with sharper boundaries than those observed in chick. Together, these results indicate conservation of the spatial organization of canonical DV and NT markers in the brown anole, with comparatively restricted expression domains at the stages examined.

We next asked whether the signaling pathways important for HAA specialization in the chick, and seen in suggestive patterns in the human, are expressed in the brown anole, and if so, in what pattern(s)? To assess this, we compared the expression of HAA-associated signaling genes across chick, human, and brown anole retinas. In the chick retina, previous studies have identified spatially restricted expression of *Fgf8, Cyp26a1, Cyp26c1*, and *Cyp1b1* (da Silva and Cepko, 2017; Joisher et al., 2025). *Bmp2* also was seen to have HAA-associated expression, although its functional role in HAA development remains untested (Joisher et al., 2025). In the human retina, spatially restricted expression patterns have been reported for retinoic acid degrading enzymes, *Cyp26a1* and *Cyp26c1* (da Silva and Cepko, 2017; Joisher et al., 2025). *Fgf8* expression has been reported to be absent from the developing human retina (da Silva and Cepko, 2017, Harding et al., 2025). Here, we examined the spatial relationships among *Cyp26a1, Cyp26c1, Bmp2*, and *Cyp1b1* in the human retina, and then asked whether corresponding expression domains were present at both developing anole foveae.

In the human retina, consistent with previous reports, *Cyp26a1* and *Cyp26c1* marked a temporal foveal domain, with *Cyp26c1* nested within the broader *Cyp26a1* territory (da Silva and Cepko, 2017, Harding et al., 2025) (Figure 2A and 2B). *Cyp1b1* showed nasal enrichment, whereas *Bmp2* was enriched in a ventral-central region, unlike the more central pattern seen in the chick (Figure 2D and 2E). Thus, as in chick, the developing human retina contains a single localized signaling domain, coincident with the developing HAA, although its molecular composition is only partially overlapping with that of chick, given the reported absence of *Fgf8* and the different expression pattern for *Bmp2* and *Cyp1b1*.

**Figure 2.**
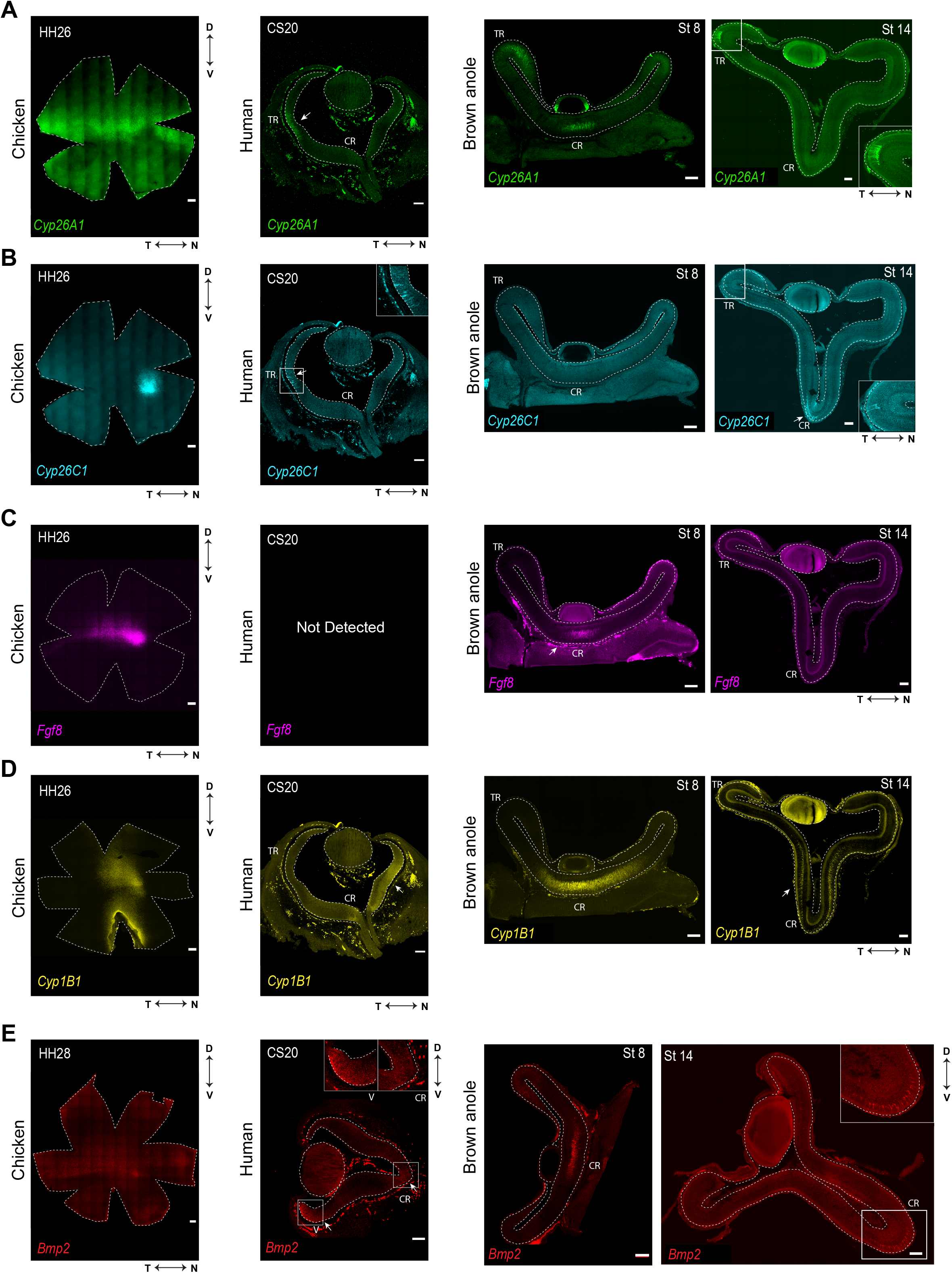
Expression of HAA-specific genes in chick, human, and brown anole retina. Comparative gene expression patterns for HAA-specific genes across developing retinas. For each panel, images are arranged from left to right as follows: chick (HH26 or HH28 as indicated); human (CS20); brown anole St8; brown anole St14. (A) *Cyp26a1;* (B) *Cyp26c1*; (C) *Fgf8;* (D) *Cyp1b1;* (E) *Bmp2*. D, dorsal; V, ventral; N, nasal; T, temporal; CR, Central Retina; TR, Temporal Retina. Scale bar: Chick (200 μm), Human (100 μm), Brown Anole (100 μm). All patterns were observed across at least 2 biological replicates (N > or = 2), except for BMP2 in the CS20 human retina, which was observed in one replicate (N=1).

In the brown anole retina, we observed two spatially distinct domains aligned with the developing central and temporal foveae. At St8, RA-degrading enzymes showed localized expression, with *Cyp26a1* present centrally and temporally, and *Cyp26c1* showing sparse central expression (Figure 2A and 2B). *Fgf8* was detected centrally and, in one sample, temporally overlapping the *Cyp26a1* and *Cyp26c1* expression domain. (Figure 2C; Supplementary Figure S2A), while *Cyp1b1* and *Bmp2* were enriched in the central retina within progenitor populations (Figure 2D and 2E). By St14, these patterns were markedly different: *Cyp26a1* became restricted to the temporal domain, *Cyp26c1* was detected centrally and sparsely in the temporal region, *Fgf8* expression was no longer detected, and *Bmp2* was expressed in a smaller central area (Figure 2, Supplementary Figure S2B-D). Together, these observations suggest that these expression domains are transient during development, while remaining spatially associated with the developing HAAs.

## Discussion

The data presented here show that a reptile shares conserved early gene expression patterns suggestive of a role in the development of a HAA. Moreover, the anole patterns suggest that the same network is used to set up two HAA’s, one in a central and one in a temporal location. It is interesting to note that most mammals do not have a HAA, and that birds, fish, and reptiles can have them, and some can have two. Mammals without a HAA nonetheless have patterned expression of the canonical DV and NT genes discussed here. A common ancestor may have deployed these genes to set up the cardinal axes. In species with a lifestyle demanding high acuity vision, perhaps including that ancestor, these axial patterning mechanisms may also have contributed to the formation of a specialized central area lacking RA signaling. Even species without a HAA, such as the mouse, have a central domain along the DV equator of *Cyp26a1* expression (Sakai et al., 2004). Early mammals, with their nocturnal lifestyle, may have lost the HAA, as it is typically important for daylight, cone-mediated vision. Some lineages leading to primates may have retained it. Alternatively, primates may have re-invented the mechanisms for HAA formation, using the DV and NT genes to position a HAA by generating a spot lacking RA signaling. RA signals are commonly used in development and patterning. Further spatial localizations and changes in the cellular composition of HAA’s may have been achieved differently in different species, with the chick using *Fgf8* to place the HAA more nasally than the human. However, the negative data on *Fgf8* expression in the human does not rule out a role for another Fgf, or even for very low expression of *Fgf8*. The anole, and perhaps other species with more than one fovea, appear to have used the RA pathway along with *Fgf8*, to set up two foveae. The RA-degrading *Cyp26a1* and *Cyp26c1* enzymes are particularly associated with the developing HAA: they are expressed nasally in the embryonic chick, temporally in the embryonic human, and both centrally and temporally in the embryonic anole (Figure 3). In contrast, *Cyp1b1*, which partially abuts the developing chick HAA, remains nasally expressed in the human retina. This may indicate that the patterning of *Cyp1b1* expression, and potentially its role in retinal development, are distinct from the other genes discussed here for HAA positioning. *Bmp2* also showed differences in spatial organization across species, with central enrichment in the anole and chick but a more ventral-central distribution in the human sample examined. Its central enrichment in the anole, rather than enrichment at both developing foveae, suggests that *Bmp2* may distinguish the molecular environments of the two foveal regions. Whether these differences contribute to foveal development remains to be determined.

**Figure 3.**
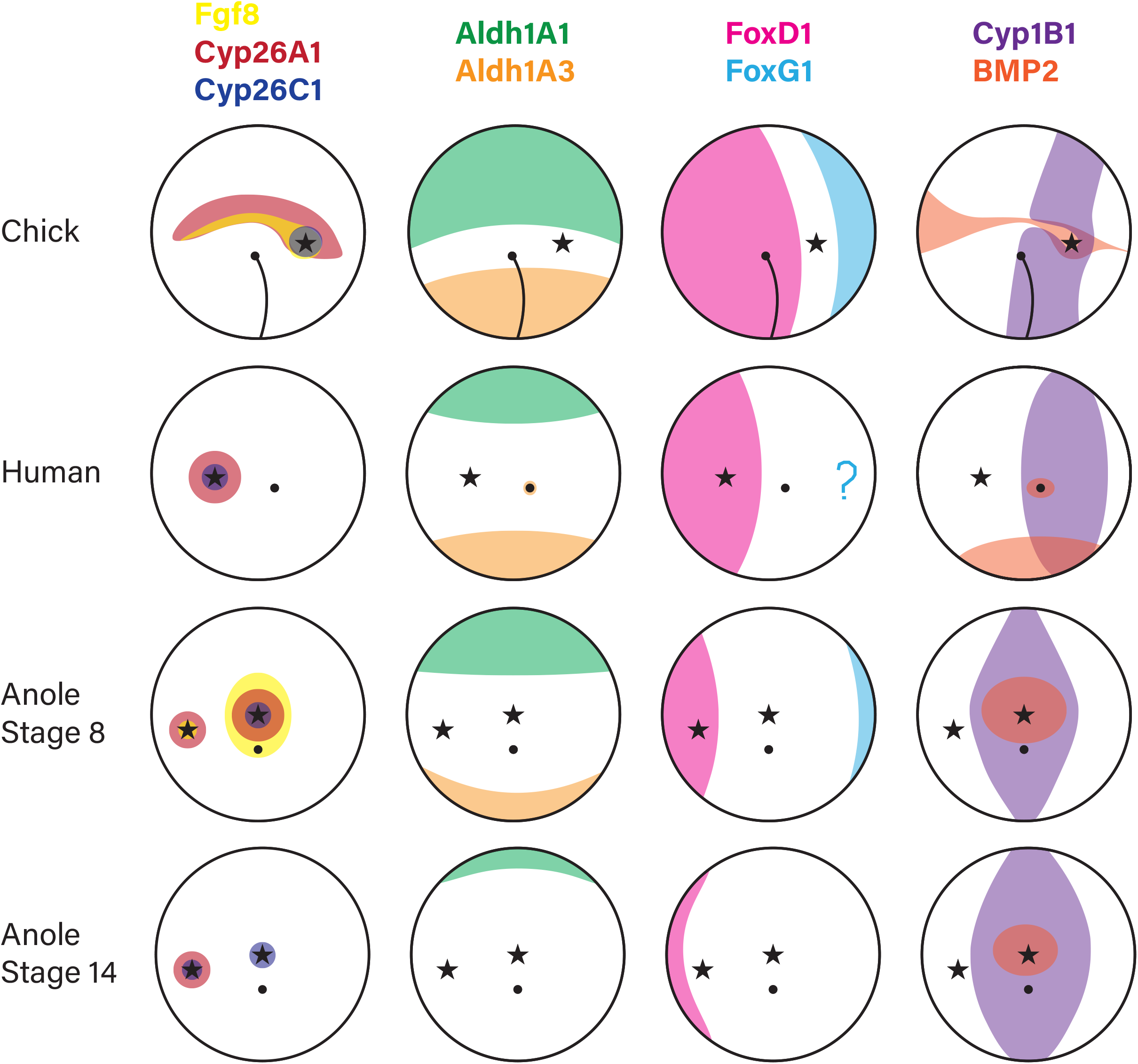
Summary of conserved DV/NT and HAA-associated expression patterns. Schematic representation of early gene expression patterns in chick (HH26/28), human (CS20), and brown anole (St8 and St14). Columns (left to right) depict expression domains of canonical DV/NT genes and genes with HAA-specific expression patterns, and the position of the HAA or fovea. Rows correspond to species and developmental stages as indicated. Black dots indicate the optic nerve head. Star symbols mark the position of HAA for each species - Chick (1 HAA: nasal-central); human (1 HAA: temporal-central), brown anole (2 HAAs: central and temporal).

In the chick retina, *Fgf8* expression and local RA degradation are required for normal HAA development: transient manipulation of RA signaling or reduction of *Fgf8* expression disrupts photoreceptor distribution, ganglion cell density, and interneuron organization (da Silva and Cepko, 2017). How these signaling domains are positioned is therefore a key question. We have proposed that the expression patterns of *Fgf8, Cyp26a1* and *Cyp26c1*, and therefore the location of the HAA, is defined by the intersecting boundaries of several early retinal axis-defining genes. We have investigated the roles of several of these axes-defining genes. Perturbation of any of three *Tbx* genes (*Tbx2, Tbx3* or *Tbx5)*, which define dorsal identity, disrupts the expression patterns of *Fgf8* and *Cyp26* genes, and alters the formation of the HAA (Ayten et al., 2026). Similarly, misexpression of *Vax*, a transcription factor that is perhaps the most upstream gene in ventral identity, perturbs the HAA rod photoreceptor pattern (Schulte et al., 2005). The *Tbx* genes and *Vax* are expressed in domains that abut the *Fgf8* and *Cyp26c1* expression spot (Joisher et al., 2025). Similar spots of *Cyp26a1*, and *Cyp26c1*, and perhaps *Fgf8*, expression mark both developing foveae in the anole.

Both anole fovea, as well as those in other species, are located equatorially along the DV axis, suggesting that conserved DV cues likely contribute to HAA positioning. In contrast, the upstream mechanisms defining the distinct NT positions of foveae, which appear to allow for the duplication of the *Fgf8* and *Cyp26* spots, remain unclear. Perhaps upstream patterning events confer competence for HAA formation in discrete NT domains for both temporal as well as central retina. This patterning may be conserved across multiple bifoveate species; bifoveate avians also generally possess a deep central fovea and secondary temporal fovea (Portier et al.,2017). These comparisons raise the possibility that early retinal pre-patterning establishes regional competence for HAA formation. It is possible that duplication of upstream NT patterning cues are present in central and temporal domains of bifoveated species.

Future gain-of-function experiments could test whether localized combinations of Fgf signaling and RA suppression are sufficient to induce a second HAA in chick or expand existing HAAs in chick or brown anole. Comparing responses across retinal locations would test whether these signals require a pre-existing permissive regional identity. Our principal comparisons focused on two developmental stages and may therefore miss intermediate changes in the onset, duration, and distribution of gene expression. Section-based analyses also limit assessment of the full extent and continuity of expression domains across the retina. Whole-mount analyses would provide a more complete spatial view, although we found that the thickness of the anole retina made flattening and imaging challenging. Tissue clearing combined with three-dimensional imaging could help overcome these limitations while preserving retinal geometry.

Together, these observations suggest that evolutionary variation in foveal number may arise through modification and redeployment of pre-existing retinal patterning programs. Further study of the anole retina could reveal novel insights into how foveal position is defined and the function of Fgf and RA pathway genes in the formation of HAAs marked by foveae.

## Acknowledgements

We thank Susana da Silva (University of Pittsburgh School of Medicine) for providing human tissue and for valuable discussions. All images were acquired using Microscopy Resources on the North Quad (MicRoN) core at Harvard Medical School. We thank Paula Montero Llopis, Praju Vikas Anekal and Adrienne Wells for their assistance with microscopy and helpful discussions.

## Materials and Methods

### Animals

Embryonic chicken retinal tissues were obtained from fertilized White Leghorn eggs (Charles River). Fertilized eggs were incubated at 38°C with humidity around 40% and embryos were staged according to Hamburger and Hamilton staging (Hamburger and Hamilton 1951). Embryonic and adult brown anole (*Anolis sagrei*) heads were shipped from University of Texas MD Anderson Cancer Center in 100% methanol on ice and stored at -20°C prior to use. Embryonic *Anolis sagrei* samples were staged as described by Sanger et al, 2008.

### Tissue preparation

#### Embryonic Chicken Tissue

Right retinas were dissected in 1X PBS and fixed overnight in 4% (v/v) paraformaldehyde (PFA) at 4°C. After fixation, retinas were washed three times for 5 minutes each in 1X PBS at room temperature (RT) before proceeding with RNA-FISH as described below. To obtain retinal cross-sections through the HAA, DV rectangles were cut out from imaged flat mounted tissue after RNA-FISH for a molecular marker for HAA (eg. *Fgf8, Cyp26c1*). The retinal pieces were then incubated in 30% sucrose in PBS (w/v) overnight at 4 °C, and transferred to a 1:1 solution (v/v) of 30% sucrose in PBS:OCT (TissueTek® O.C.T. Compound, Product code - 4583) for 3 hours. The samples were then frozen in the same 1:1 30% sucrose:OCT solution in cryomolds and stored at -80 °C. For each dorsal-HAA-ventral or nasal-HAA-temporal sample, 30 μm retina cryosections were collected and placed onto a cold superfrost plus microscope slide (Fisherbrand) within the cryostat, and stored at -80 °C.

#### Embryonic Human Tissue

All human protocols were reviewed and approved by the Institutional Review Board of Harvard. Fixed human embryonic retinal tissue from Carnegie Stages 20 and 23 was preserved in 100% methanol, and kept at -20 °C prior to use. The tissue comprised partially dissected whole eyes. To rehydrate, eyes were placed consecutively in room temperature solutions of 100%, 75%, 50%, 25% (v/v) methanol in 1X PBS, then finally into 1x PBS alone. Eyes were kept in each solution for a minimum of 15 minutes. Whole eyes were then prepared for cryosectioning in the same manner as described above. When embedding these samples, care was taken to accurately orient the tissue. At CS20, we observed a roughly pentagon-shaped pattern around the front of the eye, with one extended ventral point. This extended point was used to identify the ventral side of the retina. At CS23, some of the developing eyelid tissue was still present in the samples. The lens was closer to the nasal side of the almond-shaped opening of the developing eyelid. This feature was used to identify the nasal side of the sample.

#### Embryonic and Adult Anolis sagrei Tissue

Embryonic and adult brown anole (*Anolis sagrei*) heads were fixed overnight in 4% PFA at 4 °C. Samples were washed three times in 1× PBS (5 min each, room temperature) and dehydrated through a graded methanol series (25%, 50%, 75% methanol in 1x PBS). Heads were kept in each solution for a minimum of 15 minutes. They were next transferred to 100% methanol and shipped on ice to Harvard from University of Texas MD Anderson Cancer Center. Upon receipt, samples were stored at −20 °C in 100% methanol until further processing. Before dissections and downstream RNA-FISH, heads were rehydrated by placing them consecutively in room temperature solutions of 100%, 75%, 50%, 25% (v/v) methanol in 1X PBS, then finally into 1x PBS alone. Heads were kept in each solution for a minimum of 15 minutes. Prior to cryosectioning, heads were bisected (St 8) or whole eyes were dissected out (St 12, 14) and prepared for cryosectioning in the same manner as described above. When embedding these samples, care was taken to accurately orient the tissue.

### HCR RNA in-situ hybridization on retinal whole-mounts and cross-sections

Hybridization chain reaction (HCR) RNA in situ hybridization was performed on retinal whole-mounts of embryonic chicken and adult anole retinas using the HCR v3.0 system (Molecular Instruments) as previously described (Joisher et al., 2025). Custom probe sets were designed against target sequences (listed below) and synthesized by Molecular Instruments. Chick sequences were used for detection of anole expression patterns. Fixed retinal tissues were permeabilized in 70% ethanol diluted in PBS for 1-2 hours at room temperature. Samples were then equilibrated in wash buffer for 10 minutes at room temperature, followed by pre-hybridization in hybridization buffer at 37 °C for 30 minutes. Hybridization was carried out overnight at 37 °C in probe solution (6 nM final concentration). The following day, retinas were washed twice in pre-heated wash buffer (30 minutes each at room temperature), followed by two washes in 5× SSCT containing 0.1% Tween-20 (20 minutes each at room temperature). Samples were then equilibrated in amplification buffer for 10 minutes at room temperature. For signal amplification, matched pairs of fluorescent hairpins were snap-heated at 95 °C for 90 seconds, allowed to cool to room temperature in the dark for 30 minutes, and diluted into an amplification buffer to a final concentration of 48 nM per hairpin. Retinas were incubated in amplification solution overnight at room temperature. The next day, samples were washed twice in 5× SSCT (0.1% Tween-20) for 30 minutes each, followed by a 5 mins wash in 1x PBS and mounted for imaging using Fluoromount-G™ Mounting Medium.

HCR RNA in situ hybridization was performed on retinal cryosections from embryonic human and anole samples using the HCR protocol described above, with adjustments as previously described (Joisher et al., 2025). Slides were initially rinsed in PBS, hydrated for 5 minutes, air-dried for approximately 10 minutes (until visibly dry), and then rehydrated in PBS for an additional 5 minutes. Sections were permeabilized in 70% ethanol for 1 hour at room temperature. Following permeabilization, hybridization, amplification, and wash steps were carried out as described for whole-mount preparations. For each step, 250–300 µL of reagent was applied per slide to ensure complete coverage of the tissue section. During overnight hybridization and amplification incubations, slides were placed in humidified chambers at 37 °C and covered with hydrophobic hybridization covers to prevent evaporation.

| Species | Gene | NCBI Accession Number |
|---|---|---|
| Chicken<br>(*Gallus gallus*) | *Aldh1a1* | NM_204577.5 |
|  | *Aldh1a3* | NM_204669.3 |
|  | *Bmp2* | NM_001398170.1 |
|  | *Cyp1b1* | XM_015283751.4 |
|  | *Cyp26a1* | NM_001001129.2 |
|  | *Cyp26c1* | XM_040675448.2 |
|  | *Fgf8* | NM_001012767.2 |
|  | *FoxD1* | NM_205192.3 |
|  | *FoxG1* | NM_205193.1 |
| Brown Anole<br>(*Anolis sagrei*) | *Aldh1a1* | XM_060762194.2 |
|  | *Aldh1a3* | XM_067471297.1 |
|  | *Bmp2* | XM_060785695.2 |
|  | *Cyp1b1* | XM_060753987.2 |
|  | *Cyp26a1* | XM_060768719.2 |
|  | *Cyp26c1* | XM_067465740.1 |
|  | *Fgf8* | XM_060768258.2 |
|  | *FoxD1* | XM_067464568.1 |
|  | *FoxG1* | XM_067462863.1 |
| Human<br>(*Homo sapiens*) | *Aldh1a1* | NM_000689.5 |
|  | *Aldh1a3* | NM_000693.4 |
|  | *Bmp2* | NM_001200.4 |
|  | *Cyp1b1* | NM_000104.4 |
|  | *Cyp26a1* | NM_000783.4 |
|  | *Cyp26c1* | NM_183374.3 |
|  | *FoxD1* | NM_004472.3 |

### Imaging and Image Processing for early gene expression patterns

Samples were imaged using a Yokogawa CSU-W1 spinning disk confocal unit (50 µm pinhole size) mounted on a fully motorized Nikon Ti2 inverted microscope equipped with a motorized stage and an Andor Zyla 4.2 Plus sCMOS monochrome camera. Imaging was performed using a Plan Apo λ 20×/0.8 DIC objective and a Plan Fluor 40×/1.3 Oil DIC objective. Image acquisition was controlled using Nikon NIS-Elements AR (v5.02).

Z-stacks were collected using a Piezo Z-drive, with the excitation shutter closed during axial movement to minimize photobleaching. Raw image data were saved in ND2 format. Image analysis was performed in FIJI (ImageJ). For each sample, maximum intensity projections were generated from z-planes containing robust in situ hybridization signal. Brightness and contrast adjustments were applied uniformly across comparable samples to improve visualization of signal.

## Supplementary Figure Legends

**Figure S1. *FoxG1* expression in developing chick and brown anole retina**.

Comparative gene expression patterns for *FoxG1* across developing chicken and anole retinas. For each panel, images are arranged from left to right as follows: chick (HH26); brown anole St8; brown anole St14. D, dorsal; V, ventral; N, nasal; T, temporal; CR, Central Retina; TR, Temporal Retina. Scale bar: Chick (200 μm), Brown Anole (100 μm). All patterns were observed across at least 2 biological replicates (N > or = 2).

**Figure S2. Fgf8 and RA-associated gene expression patterns in developing brown anole retina**.

**(A)** *Fgf8* expression in NT retinal cross-section at St8. **(B)** *Fgf8* expression in DV retinal cross-section at St8, St12 and St14. **(C)** *Cyp26a1* expression in DV retinal cross-section at St8 and St14. **(D)** *Cyp26c1* expression in DV retinal cross-section at St8, St12 and St14. Insets show higher-magnification views of boxed regions. The tissue that appears to be present inside the eye in several samples is the result of deflation of the eye during storage in methanol, causing sections of the retina to dent inward and be sectioned at an oblique angle. D, dorsal; V, ventral; T, temporal; N, nasal; CR, Central Retina; TR, Temporal Retina. Scale bar: 100 μm. Pattern in A was observed in one sample (N=1); patterns in B-D were each observed in two samples (N=2), except St12 *Fgf8* which was observed in one sample (N=1).

## References

1. Sanger TJ, Losos JB, Gibson-Brown JJ. A developmental staging series for the lizard genus Anolis: a new system for the integration of evolution, development, and ecology. J Morphol. 2008; 269(2): 129–137. 10.1002/jmor.10563.

2. Hamburger, V. and Hamilton, H.L., 1951. A series of normal stages in the development of the chick embryo. Journal of morphology, 88(1), pp.49–92.

3. Harding, P., Wojtynska, M., Smith, A.J., Ali, R.R. and Pearson, R.A., 2025. Human macula formation involves two waves of retinoic acid suppression via CYP26A1 that modulate cell cycle exit and cone subtype specification. Stem Cell Reports, 20(10).

4. Joisher, H.N., Lee, C., Prabhakara, C., van der Weide, I., Si, Y., Lonfat, N. and Cepko, C., 2026. Integration of in situ hybridization and scRNA-seq data provides a 2D topographical map of the developing retina across species. bioRxiv, pp.2026–01.

5. da Silva, S. and Cepko, C.L., 2017. Fgf8 expression and degradation of retinoic acid are required for patterning a high-acuity area in the retina. Developmental cell, 42(1), pp.68–81.

6. Takahashi, H., Sakuta, H., Shintani, T. and Noda, M., 2009. Functional mode of FoxD1/CBF2 for the establishment of temporal retinal specificity in the developing chicken retina. Developmental biology, 331(2), pp.300–310.

7. Sakuta, H., Takahashi, H., Shintani, T., Etani, K., Aoshima, A. and Noda, M., 2006. Role of bone morphogenic protein 2 in retinal patterning and retinotectal projection. Journal of Neuroscience, 26(42), pp.10868–10878

8. O’Brien, J.J., Chen, X., MacLeish, P.R., O’Brien, J. and Massey, S.C., 2012. Photoreceptor coupling mediated by connexin36 in the primate retina. Journal of Neuroscience, 32(13), pp.4675–4687.

9. Dütting, D. and Thanos, S., 1995. Early determination of nasal-temporal retinotopic specificity in the eye anlage of the chicken embryo. Developmental biology, 167(1), pp.263–281. 19.

10. Echelard, Y., Epstein, D.J., St-Jacques, B., Shen, L., Mohler, J., McMahon, J.A. and McMahon, A.P., 1993. Sonic hedgehog, a member of a family of putative signaling molecules, is implicated in the regulation of CNS polarity. Cell, 75(7), pp.1417–1430.

11. Crossley, P.H., Martinez, S. and Martin, G.R., 1996. Midbrain development induced by FGF8 in the chicken embryo. Nature, 380(6569), pp.66–68.

12. Matsuno, T., Itasaki, N., Ichijo, H. and Nakamura, H., 1992. Retinotectal projection after partial ablation of chicken optic vesicles. Neuroscience research, 15(1-2), pp.96–101.

13. McLaughlin, T. and O’Leary, D.D., 2005. Molecular gradients and development of retinotopic maps. Annu. Rev. Neurosci., 28(1), pp.327–355.

14. Hendrickson A: A morphological comparison of foveal development in man and monkey. Eye 1992, 6:136–144

15. Hendrickson, A.E. and Yuodelis, C., 1984. The morphological development of the human fovea. Ophthalmology, 91(6), pp.603–612.

16. Sugiyama, T., Yamamoto, H., Kon, T., Chaya, T., Omori, Y., Suzuki, Y., Abe, K., Watanabe, D. and Furukawa, T., 2020. The potential role of Arhgef33 RhoGEF in foveal development in the zebra finch retina. Scientific Reports, 10(1), p.21450.

17. Rasys, A.M., Pau, S.H., Irwin, K.E., Luo, S., Kim, H.Q., Wahle, M.A., Trainor, P.A., Menke, D.B. and Lauderdale, J.D., 2021. Ocular elongation and retraction in foveated reptiles. Developmental Dynamics, 250(11), pp.1584–1599.

18. Sannan, N.S., Shan, X., Gregory-Evans, K., Kusumi, K. and Gregory-Evans, C.Y., 2018. Anolis carolinensis as a model to understand the molecular and cellular basis of foveal development. Experimental Eye Research, 173, pp.138–147.

19. Potier, S., Mitkus, M., Bonadonna, F., Duriez, O., Isard, P.F., Dulaurent, T., Mentek, M., Kelber, A., 2017. Eye size, fovea, and Foraging ecology in Accipitriform raptors. Brain Behav. Evol. 90, 232–242.

20. Rasys, A.M., Wegerski, A., Trainor, P.A., Hufnagel, R.B., Menke, D.B. and Lauderdale, J.D., 2024. Dynamic changes in ocular shape during human development and its implications for retina fovea formation. BioEssays, 46(1), p.2300054.

21. Rasys, A.M., Pau, S.H., Irwin, K.E., Luo, S., Kim, H.Q., Wahle, M.A., Menke, D.B. and Lauderdale, J.D., 2025. Histological analysis of retinal development and remodeling in the brown anole lizard (Anolis sagrei). Journal of Anatomy, 246(6), pp.1019–1033.

22. Wahle, M.A., Kim, H.Q., Menke, D.B., Lauderdale, J.D. and Rasys, A.M., 2023. Maturation and refinement of the maculae and foveae in the Anolis sagrei lizard. Experimental eye research, 234, p.109611.

23. Ayten, M., Joisher, H. N. V., Prabhakara, C., & Cepko, C. (2026). Tbx genes influence early gene expression and photoreceptor patterning in the chick retina. Developmental Biology, 533, 61–74.

24. Pittlik, S., Domingues, S., Meyer, A., & Begemann, G. (2008). Expression of zebrafish aldh1a3 (raldh3) and absence of aldh1a1 in teleosts. Gene Expression Patterns, 8(3), 141–147.

25. Fan, X., Molotkov, A., Manabe, S., Donmoyer, C. M., Deltour, L., Foglio, M. H., Cuenca, A. E., Blaner, W. S., Lipton, S. A., & Duester, G. (2003). Targeted disruption of Aldh1a1 (Raldh1) provides evidence for a complex mechanism of retinoic acid synthesis in the developing retina. Molecular and Cellular Biology, 23(13), 4637–4648.

26. Sakai, Y., Luo, T., McCaffery, P., Hamada, H. and Dräger, U.C., 2004. CYP26A1 and CYP26C1 cooperate in degrading retinoic acid within the equatorial retina during later eye development. Developmental Biology, 276(1), pp.143–157.

27. Schulte, D., Peters, M.A., Sen, J. and Cepko, C.L. (2005). The rod photoreceptor pattern is set at the optic vesicle stage and requires spatially restricted cVax expression. Journal of Neuroscience, 25(11), 2823–2831.

