## Supplemental Figure 1 and 2 for "Gene expression in *Anolis sagrei* suggests conservation of patterning mechanisms drives formation of two high acuity areas"

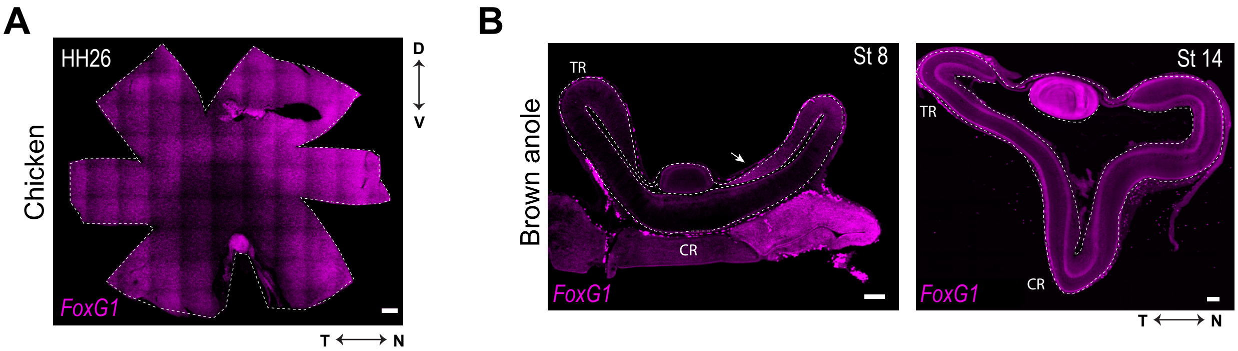

A
HH26
Chicken
D
V
FoxG1
T
N
B
Brown anole
St 8
TR
CR
FoxG1
St 14
TR
CR
FoxG1
T
N

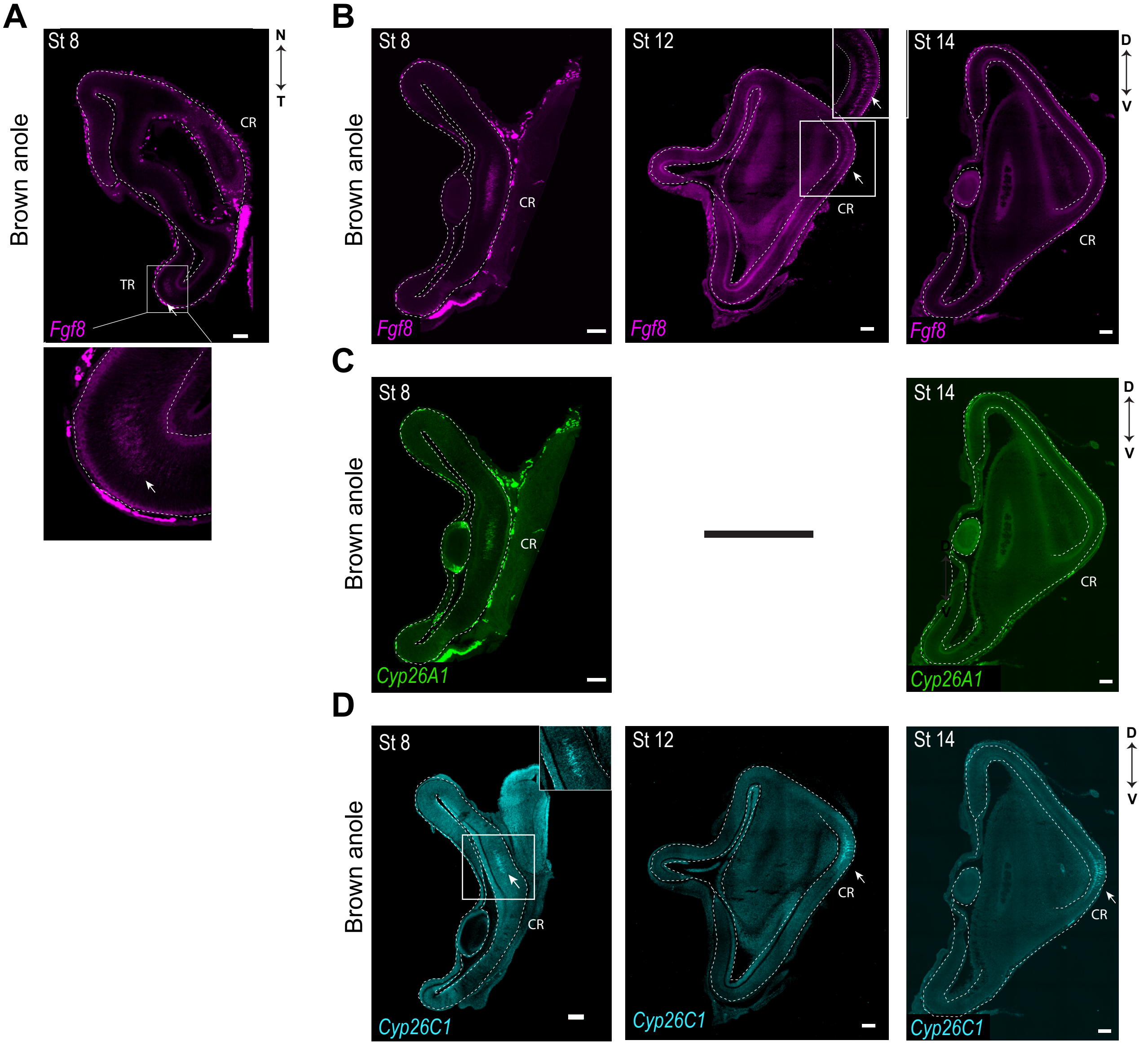
A
B
C
D
Brown anole
St 8
St 12
St 14
N
T
D
V
CR
TR
Fgf8
Cyp26A1
Cyp26C1
